# Empathi: Embedding-based Phage Protein Annotation Tool by Hierarchical Assignment

**DOI:** 10.1101/2024.12.31.630607

**Authors:** Alexandre Boulay, Audrey Leprince, François Enault, Elsa Rousseau, Clovis Galiez

## Abstract

Bacteriophages, viruses infecting bacteria, are estimated to outnumber their cellular hosts by 10-fold, acting as key players in all microbial ecosystems. Under evolutionary pressure by their host, they evolve rapidly and encode a large diversity of protein sequences. Consequently, the majority of functions carried by phage proteins remain elusive. Current tools to comprehensively identify phage protein functions from their sequence either lack sensitivity (those relying on homology for instance) or specificity (assigning a single coarse grain function to a protein). Here, we introduce Empathi, a protein-embedding-based classifier that assigns functions in a hierarchical manner – from general functional categories such as “structural” and “DNA-associated” proteins to more specific ones including “nucleases”, “tail appendages” and “endolysins” to name only a few. These categories were specifically tailored for phage protein functions and organized such that molecular-level functions are respected in each category, making it well suited for training machine learning classifiers based on protein embeddings. We show on a dataset of cultured phage genomes that Empathi significantly outperforms homology-based methods, tripling the number of annotated homologous groups. On the EnVhog database, the most recent and extensive database of metagenomically-sourced phage proteins, Empathi doubled the annotated fraction of protein families from 16% to 33%. On complete genomes taken from new viromes, almost twice as many proteins are annotated using our method, predictions are consistent when compared to existing tools and Empathi predictions are highly colocalized. In addition, by leveraging Empathi’s ability to assign multiple labels to the same protein, it is possible to identify multifunctional proteins such as virion-associated lysins. Having a more global view of the repertoire of functions a phage possesses will assuredly help to understand them and their interactions with bacteria better.

## 1 Introduction

Bacteriophages or phages – viruses that infect bacteria – are some of the most abundant biological entities on earth, being present everywhere from the ocean and the soil to our very own bodies^1,2^. Despite this, phages have been overlooked in studies of the microbiome that mainly focus on the bacterial component. Furthermore, wet-lab studies on phages are slow and labor intensive, traditionally requiring phages to be cultured in the presence of a known bacterial host. The recent development of next-generation sequencing methods, especially shotgun sequencing, has substantially accelerated the study of phages, allowing to sequence them directly in their natural habitats, thereby circumventing phage culturing. However, being able to assemble new phage genomes from massive metagenomics sequencing data comes with the challenge of characterizing these highly diverse phages and the proteins they are composed of, as well as determining their host.

Phage protein function prediction is a major challenge; it was recently shown by Pérez-Bucio et al^3^ that only 16% of the diversity of phage protein families has been assigned a function. This issue has been the focus of numerous studies^4–22^, most limiting their efforts to predicting one type of protein at a time. The most studied of these are phage virion proteins (PVPs), the structural proteins of phages^4–16,23–26^. In particular, receptor binding proteins (RBPs) that allow binding and recognition of the bacterial host^17,18^, and lysins that degrade the peptidoglycan or exopolysaccharide layers surrounding and protecting bacteria^19–22^ have received significant research attention.

Currently, the most widely used method for assigning general functional annotations to phage proteins is an approach based on profile hidden Markov models (pHMMs) which relies on sequence homology^27^. Yet, as described by Flamholz *et al.*^28^, in viral metagenomics, these methods are constrained by the limited number of annotated proteins that can be used to construct probabilistic sequence models and by the high mutation rate of viruses resulting in a large diversity of protein sequences.

Recently, alignment-free methods based on protein embeddings, fixed-size real-valued vectors obtained from protein language models (PLMs) such as ProtTrans^29^ and ESM2^30^, were developed and leverage functional homology between proteins. These models have been trained on an enormous corpus of protein sequences and have been demonstrated to capture the structural and functional information from protein sequences^28,31^.

Different tools based on protein embeddings were recently developed for phage protein function annotation^5,18,19,22,28^, with all but one being specific to particular protein families. For example, Yang *et al*.^18^ trained a classifier to identify tail-spike proteins with a distinctive beta-helix domain, and Concha-Eloko *et al.* proposed DepoScope^22^, a tool trained for depolymerase detection and functional domain identification. The only classifier able to predict general phage protein functions, VPF-PLM^28^, is a multilabel classifier that uses embeddings obtained using ESM2 and that is trained to predict the basic PHROG categories^32^. However, these categories possess overlapping molecular-level functions creating noise in the training data that hinders the accuracy and sensitivity that can be achieved by models trained on them.

In this paper we present Empathi, an **Em**bedding-based **P**hage **P**rotein **A**nnotation **T**ool that **Hi**erarchically assigns proteins functions. To this end, a hierarchical scheme for functional categories that respects molecular-level functions was defined from the PHROG classification^32^ to be better adapted for machine learning (ML) classification. Empathi is composed of multiple binary models trained on proteins from completely sequenced phages. These models are then used to assign functions to new proteins, starting from general annotations such as “structural” or “DNA-associated” to more precise functions when possible. Empathi significantly outperforms homology-based methods, tripling the number of annotated homologous groups in our original dataset of cultured phage genomes. Applying it to the EnVhog^3^ database of metagenomically sourced phage proteins, Empathi more than doubled the number of previously annotated protein families from 16% to 33%. Finally, on complete genomes taken from new viromes, almost twice as many proteins are annotated using our method and predictions are consistent when compared to existing tools.

## 2 Methods

### 2.1 Defining a new hierarchical scheme for functional annotations of phage proteins

PHROG categories are not adapted for machine learning classification. Indeed, many PHROG categories encompass various molecular-level functions. For example, the head and packaging category is composed of structural proteins, internal proteins with lytic domains, terminase proteins that can bind DNA, etc. Another example are DNA-associated proteins that can be found in 7 out of the 9 PHROG categories.

Here, from a biological perspective and with machine learning purposes in mind, similar PHROG annotation terms were grouped together into new functional categories that respect molecular functions (see Figure 1). The complete list of PHROG annotation terms constituting each newly defined functional category can be found in the repository in *data/functional_groups.json*. These include groups such as “baseplate proteins”, “nucleases” and “adsorption-related proteins”. These newly defined functional categories are, when possible, classified into more general ones (“PVP”, “DNA-associated”, “lysis-associated”). Proteins can be associated with multiple categories. For example, tail proteins with lytic activity are included in the PVP category and in the “cell wall depolymerase” category.

**Figure 1.**
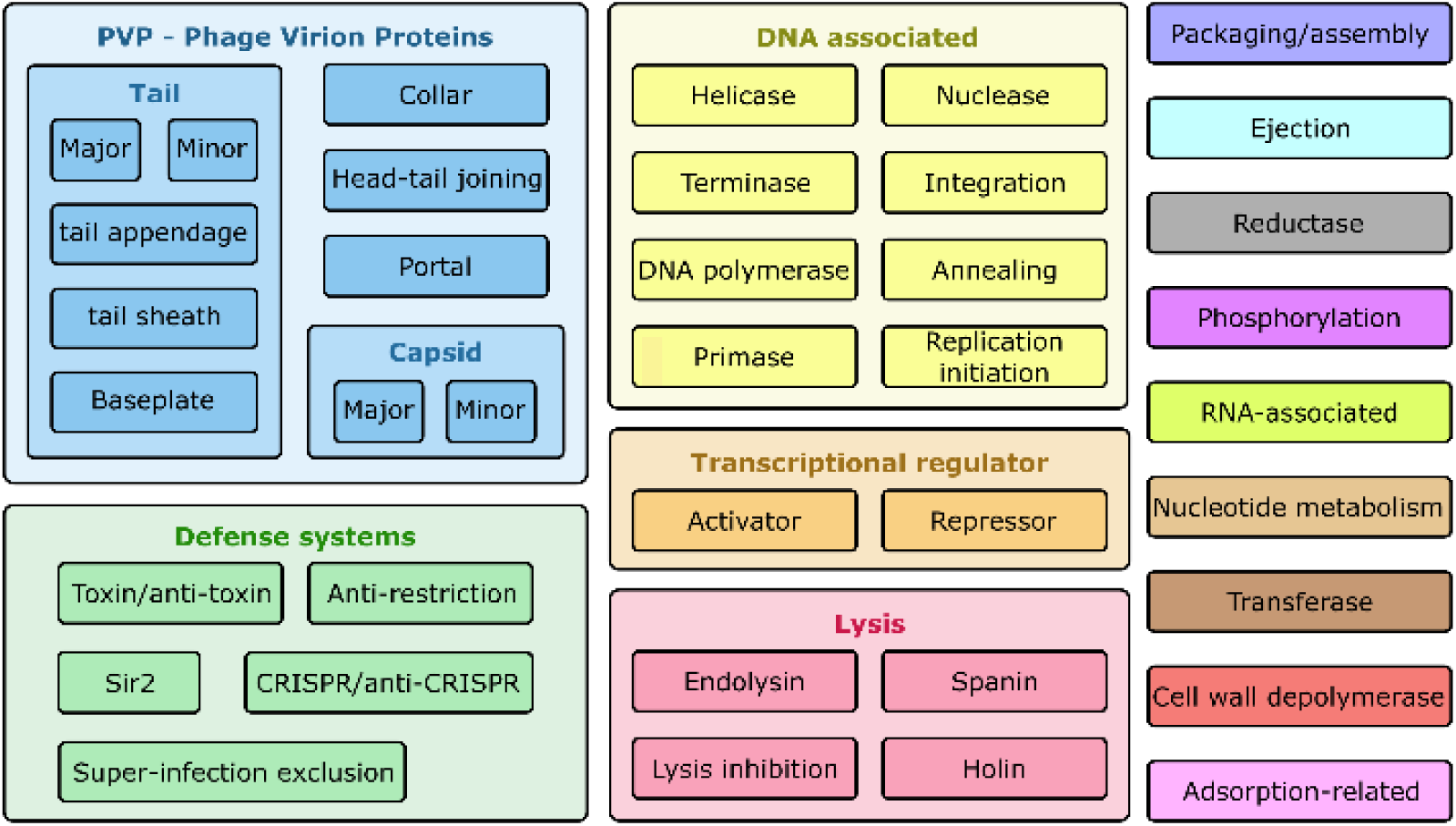
Definition of hierarchical functional categories used in Empathi. PHROG annotation terms are placed in all categories they fit in (e.g. tail protein with lytic activity are placed in “tail”, “pvp” and “cell wall depolymerase”). The colours defined here for each category were used in all following figures.

### 2.2 Collecting and annotating phage genomes and proteins

Using INPHARED^33^, 18,477 phage genomes with were collected from GenBank^34^ (Figure 2) along with their predicted protein sequences on January 2^nd^ 2024. A total of 1.85M phage proteins predicted by Prokka^35^ (which employs Prodigal^36^) were deduplicated (identity of 100%) into 904k unique proteins. To annotate them functionally, these proteins were compared with the pHMMs of the PHROG^32^ database using HH-suite^37^, the best hit being considered for each protein (e-value less than 0.001). Using ProtTrans^29^, a protein language model learned on billions of tokens (amino acids), embeddings in the form of fixed-size 1024-dimensional vectors were computed for every protein in the dataset. Finally, proteins were clustered using MMseqs2^38^ using a 30% sequence identity threshold, 80% coverage and an e-value less than 0.001 to create training and testing sets for machine learning.

**Figure 2.**
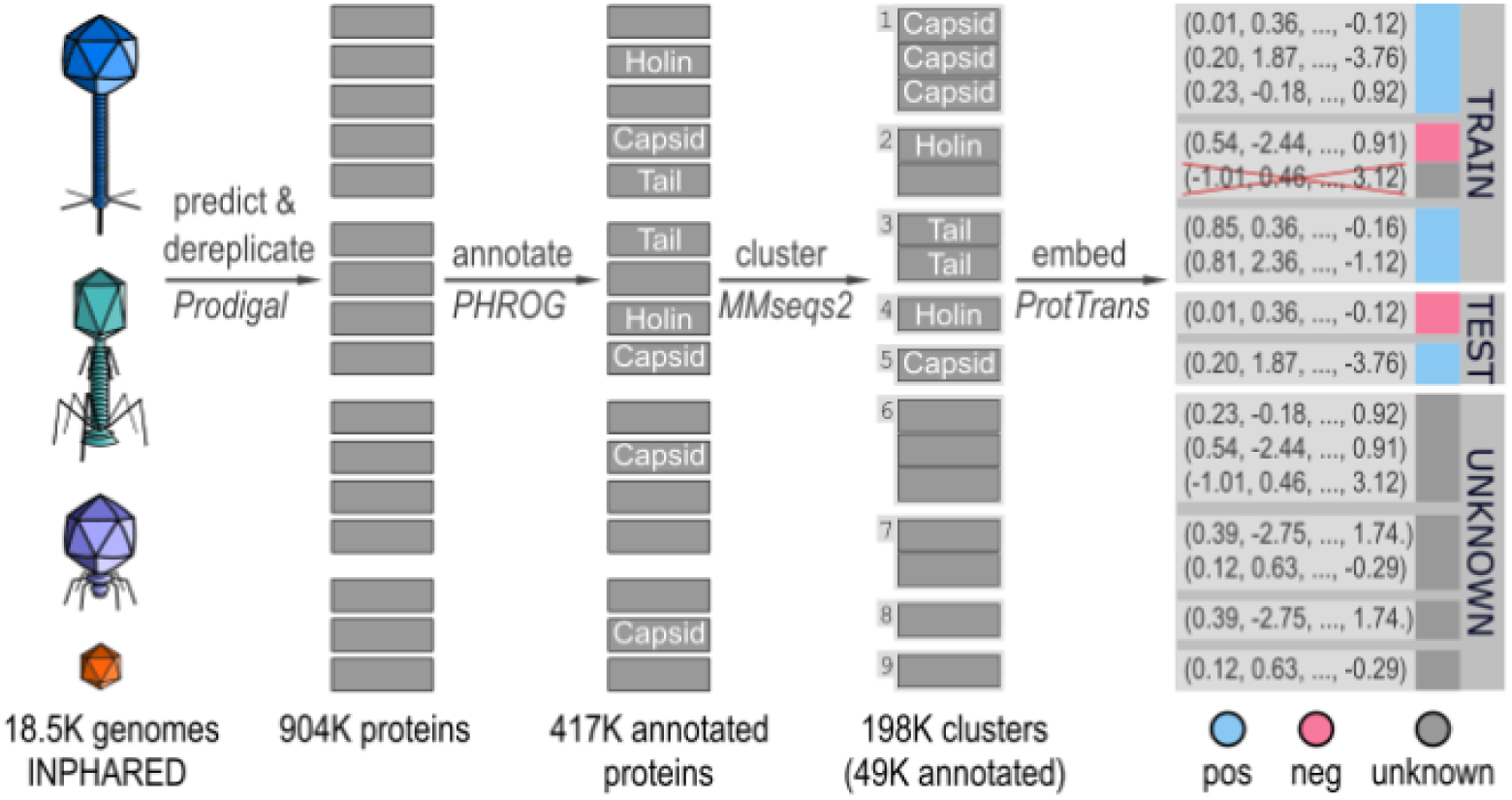
Dataset preparation for machine learning. Phage genomes were downloaded from public databases using INPHARED. Protein sequences were predicted using Prodigal and annotated by sequence-HMM comparison with proteins of known function in the PHROG database. Proteins were clustered using MMseqs2 and embedded using ProtTrans. To train models, all proteins with known functions from one cluster are either placed in the training set or in the testing set. The positive and negative labels are defined independently for each binary model that is trained; in this example, the positive set is composed of phage virion (structural) proteins.

### 2.3 Defining training and testing sets

For each functional category in Figure 1, a subset of the dataset was constituted, containing positive and negative examples of that category (Supplementary table 1). Only clusters containing proteins with no contradictory annotations (associated to the positive and negative set) were included. Proteins with unknown functions, even if they are found in a cluster with other proteins with known functions, are not considered when training models. To constitute the positive class, all proteins assigned to this category were considered. Both to ensure a diverse selection of proteins in the negative class and to make positive and negative classes as homogenous as possible in size, only one protein per MMseqs2 cluster was incorporated in the negative class for general categories. For sub-categories, only proteins from the parent category are considered. Consequently, less proteins are available to constitute the negative set and therefore all proteins from the negative class (all proteins per cluster) were considered when training these models. For most functional categories, the negative class still contained 3-4 times more proteins than the positive one (Supplementary table 1). To correct for this imbalance, weights on model per-class performance (1 / frequency of each class) were added during training. In some cases, proteins with very general annotations were left out of both positive and negative classes. For example, when training a model to predict tail proteins, proteins only annotated as “phage virion proteins” were excluded as these could correspond to tail or non-tail proteins. Finally, to limit data leakage between training and testing sets, all proteins from one MMseqs2 cluster were either included in the training or in the testing set. For each category, the training set is built by randomly sampling 80% of clusters, leaving the remaining 20% to constitute the testing set.

### 2.4 Training and testing models

Empathi is trained on the annotated portion of phage proteins obtained from INPHARED. Support vector machines (SVM) with an RBF (Radial Basis Function) kernel were used as the base classifier in all models.

For each functional category shown in figure 1 and their associated positive and negative classes, a binary model was trained on the corresponding training set as explained in the previous section. In this way, the classifier for each category was both trained and tested independently from all other categories.

A hierarchical approach was implemented (Figure 1): models for sub-categories were trained using only the proteins in the parent category. This means for example, that a protein must first be predicted as being a “tail protein” before it is assigned (or not) the “baseplate” annotation. As more data is used to train the parent classes, this approach results in higher confidences and recall on general functions while allowing to assign specific functions when possible. Furthermore, this approach reduces the required computational resources as models of subcategories are only applied when their parent category was assigned.

In addition, having models that are binary and hierarchical makes them accurate for all functional categories despite some proteins corresponding to several groups. Returning to a previous example, a protein can thus be classified as being a tail protein (“tail” and “PVP”) that recognizes a bacterial receptor (“adsorption-related”) and/or degrades the bacterial cell wall (“cell wall depolymerase”).

F1-score, precision and recall were computed to measure the performance of each binary model and were reported as a function of the confidence of predictions.

### 2.5 Additional evaluation of robustness by better separating remote homologs

Next, we wanted to evaluate the robustness of our approach, in particular, to see how well our model can generalize to proteins that are very different from those seen during training. To this end, proteins from a given homologous group (a PHROG) and all proteins from similar PHROGs were removed from the training set (similarities between PHROGs being indicated in the PHROG database). Then, a model was trained, excluding these proteins, and its performance evaluated on the holdout PHROGs. As a new model needs to be retrained for every iteration, this procedure is computationally intensive and was only performed ten times for six chosen categories (“lysis”, “cell wall depolymerase”, “DNA-associated”, “PVP”, “transcriptional regulator”, “transferase”).

### 2.6 Applying Empathi to the EnVhog database and to complete phage genomes from new viromes

Firstly, three metaviromes were obtained from Garmaeva *et al*.^39^ (gut), Bi *et al*.^40^ (sulfuric soil), and Ni *et al*.^41^ (marine water). Complete genomes were identified using CheckV^42^ and two genomes were randomly chosen from each study. Proteins were predicted using Prodigal, embedded using ProtTrans and Empathi was used to predict their functions. Predictions were compared to the annotations obtained by PHROG pHMMs (HH-suite; best hit with e-value < 1e^-3^) and to the predictions made by VPF-PLM using their default thresholds.

Secondly, the EnVhog^3^ database composed of phage proteins obtained from metagenomic experiments was downloaded (on June 20^th^ 2024) and protein functions were predicted with Empathi. No additional manipulations were required as protein embeddings for representative sequences of each EnVhog cluster together with their corresponding PHROG annotations were available. EnVhog clusters are created as described in Pérez-Bucio *et al.* by using profile HMMs of sequence-level clusters at 30% identity.

## 3 Results

Here, we present Empathi, a tool to functionally annotate phage proteins. First, the methodology was evaluated on a set of phage proteins with known functions and then employed to tentatively assign a function to proteins with no hits to proteins with known functions. Next, we demonstrate that our approach can be used on new phage genomes (from new viromes), showing consistency of predictions through genomic maps that illustrate the colocalization of protein functions. We also used Empathi to annotate the EnVhog database, more than doubling the number of functionally annotated proteins compared to pHMM annotation. Throughout all these analyses, we compare our performances to state-of-the-art approaches such as pHMM homology identification and VPF-PLM^28^.

### 3.1 Improved precision and recall for viral protein annotation

#### Building and testing models for all newly defined functional categories

Among the 904k dereplicated proteins from INPHARED genomes, 417k (46%) were placed in at least one of the 44 newly defined functional categories (Figure 1) based on their sequence similarity to PHROG HHMs. Of the 198k clusters generated using MMseqs2, 49k (25%) contained annotated proteins and their structure was used to separate proteins in the training and testing sets in order to reduce data leakage due to homology.

A binary model was trained for each of the 44 functional categories using 80% of clusters and tested on the remaining 20% of clusters (Supplementary table 1). The F1-score for all binary models except one was greater than or equal to 88% with three quarters of models reaching scores of at least 95% (Supplementary table 1). Only the model trained to predict collar proteins was less performant, achieving a score of 60% (Supplementary figure and table 1). This is most likely due to the very limited number of collar proteins in the dataset (233 proteins corresponding to 39 clusters in the training set). The precision and recall curves demonstrate that models are confident in their predictions, being able to achieve an excellent recall (83-100% at 50% confidence) whilst conserving an excellent precision (80-100% at 50% confidence) (Figure 3A, supplementary figure and table 1).

**Figure 3.**
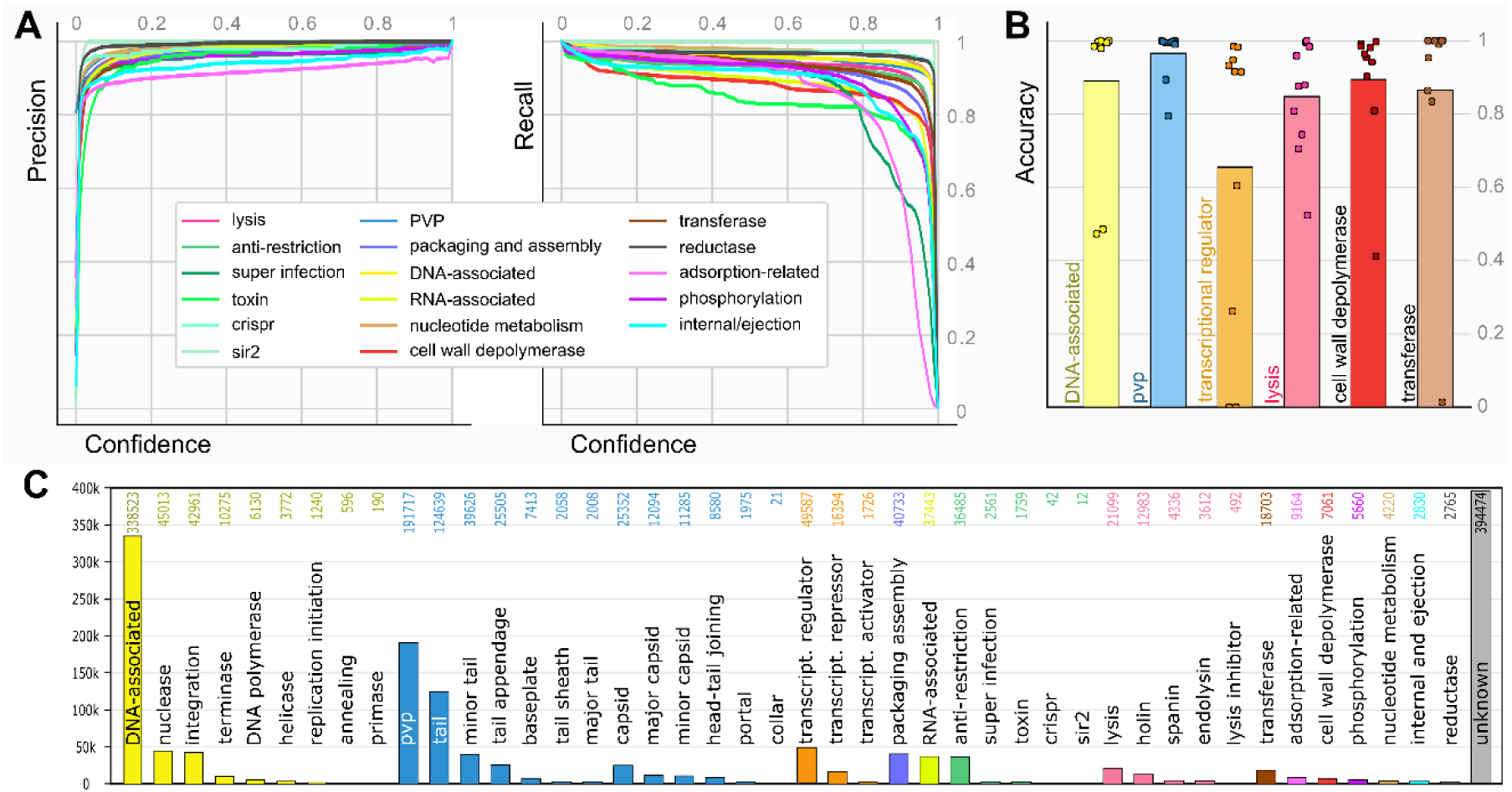
Evaluation of the performance and robustness of Empathi, and its predictions on previously unannotated proteins. **A)** Precision and recall curves as a function of confidence (prediction probability) for Empathi binary models trained on the general functional groups. PVP: Phage virion protein. **B)** Robustness analysis: accuracy of models on holdout PHROGs. Bars represent the average performance over 10 iterations. **C)** Predicted protein counts for each functional group in the fraction of the original dataset (INPHARED) that did not align to proteins with known functions in the PHROG database.

Many of the errors made by the models stem from biologically similar functions. The baseplate model classified tail appendage proteins as baseplate proteins and vice versa reflecting the biological similarity of these proteins. Similarly, the primase model incorrectly classified some helicases. In addition, some proteins annotated only as *tail proteins* are predicted by our model as being adsorption-related proteins. These are likely not errors, but the protein annotations are too general to validate our predictions.

Our tool’s consistency was tested by looking at the predictions made within each cluster of homologous proteins. Because these proteins are very similar, the predictions they receive from our model should also be similar. When applying Empathi to the proteins of the 197,739 clusters in the whole dataset (training set + testing set + unknown), 163,569 clusters (83%) indeed received identical predictions for every protein. 11,361 clusters (6%) received identical predictions with some proteins receiving no function. The proteins in 16,418 clusters (8%) were predicted as sharing at least one general function but were assigned either differing specific functions or a differing second general function. Finally, only 6,391 clusters (3%) received differing predictions.

#### Evaluation of robustness

Even though proteins detected as similar by sequence-sequence comparisons were separated into either the training or the testing set, remote homologs whose similarity is not detected by MMseqs2 might be present in both sets. As HMM-HMM comparisons of the PHROGs detect more distant homologous relationships between proteins, new training sets were built by removing one PHROG as well as any PHROG similar to it. In this manner, retrained models were tested on proteins that had no resemblance, even remote, to any protein in the training dataset.

Overall, models predicted the function of interest accurately for these holdout PHROGs (Figure 3B), having an accuracy greater than 70% in 51 of the 60 replicates (85%). Lower model performances on some holdout groups correspond to cases where the holdout group was very large and where all examples of a specific type of protein (e.g. MotB-like transcriptional regulator) have been removed from training by the holdout procedure.

#### Expanding the proportion of phage proteins with annotated functions

As mentioned previously, as many as 487k proteins (54%) corresponding to 150k clusters (75%) were not similar to any PHROG pHMM with a known function. Considering these proteins that even sensitive similarity search methods have failed to functionally annotate, 62% (300k proteins) were assigned a function with Empathi. In total, almost four phage proteins out of five now have a predicted function (718k proteins, 79% of the dataset), bringing the total ratio of annotated clusters from 25% to 73.5%. On average, the confidence of the highest-level prediction for previously unannotated proteins is 80.3%.

A high diversity of protein functions is observed in this fraction of the dataset (Figure 3C). “DNA-associated” (33%) and “phage virion proteins” (20%) are the functions that were assigned to the most proteins. Among the predicted PVPs, about two thirds are tail proteins. In addition, 49.5k transcriptional regulators, 45k nucleases, 43k integration-related, 40.7k packaging/assembly related, 37.4k RNA-associated and 36.5k anti-restriction proteins were also identified, highlighting the unexplored diversity of these proteins.

### 3.2 Increased number and high colocalization of protein functions assigned to complete genomes from new viromes

Complete phage genomes were obtained from three viromes^39–41^ sampled from various environments (human gut, sulfuric soil and marine water) and, for the purpose of this analysis, two complete genomes were randomly picked from each study. Four of these aligned to *Caudoviricetes* species using a BLAST^43^ search of the core nucleotide database: LN_7A03 at 92% identity and 36% coverage, LN_4A01 at 88% identity with 8% query coverage, and both PP079085.1 and PP079056.1 at 100% identity with 100% query coverage (see supplementary data for their genomic sequences). Both sequences from the Bi *et al.* study on sulfuric soils, M8_k141 and M4_k141 did not return any significant hits using BLAST.

Functions were assigned using Empathi, VPF-PLM and PHROG pHMMs to the proteins from these genomes. A greater number of proteins were assigned a function using Empathi than the two other approaches. Out of a total of 574 proteins for the six genomes presented in Figure 4, 414 proteins were assigned a function using Empathi, compared to 232 using the PHROG database and 202 using VPF-PLM. For the most part, the annotations obtained from all three methods are coherent. For example, most “transcriptional regulators”, “integration proteins”, “RNA-associated” and “nucleotide metabolism” proteins from Empathi categories are consistent with the “DNA, RNA and nucleotide metabolism” PHROG category, thus colored in yellow-orange in both cases. The same is true for “phage virion proteins” (PVPs), “packaging proteins” and “internal proteins” from Empathi categories being consistent with the “tail”, “head and packaging” and “connector” PHROG categories depicted in varying shades of blue.

**Figure 4.**
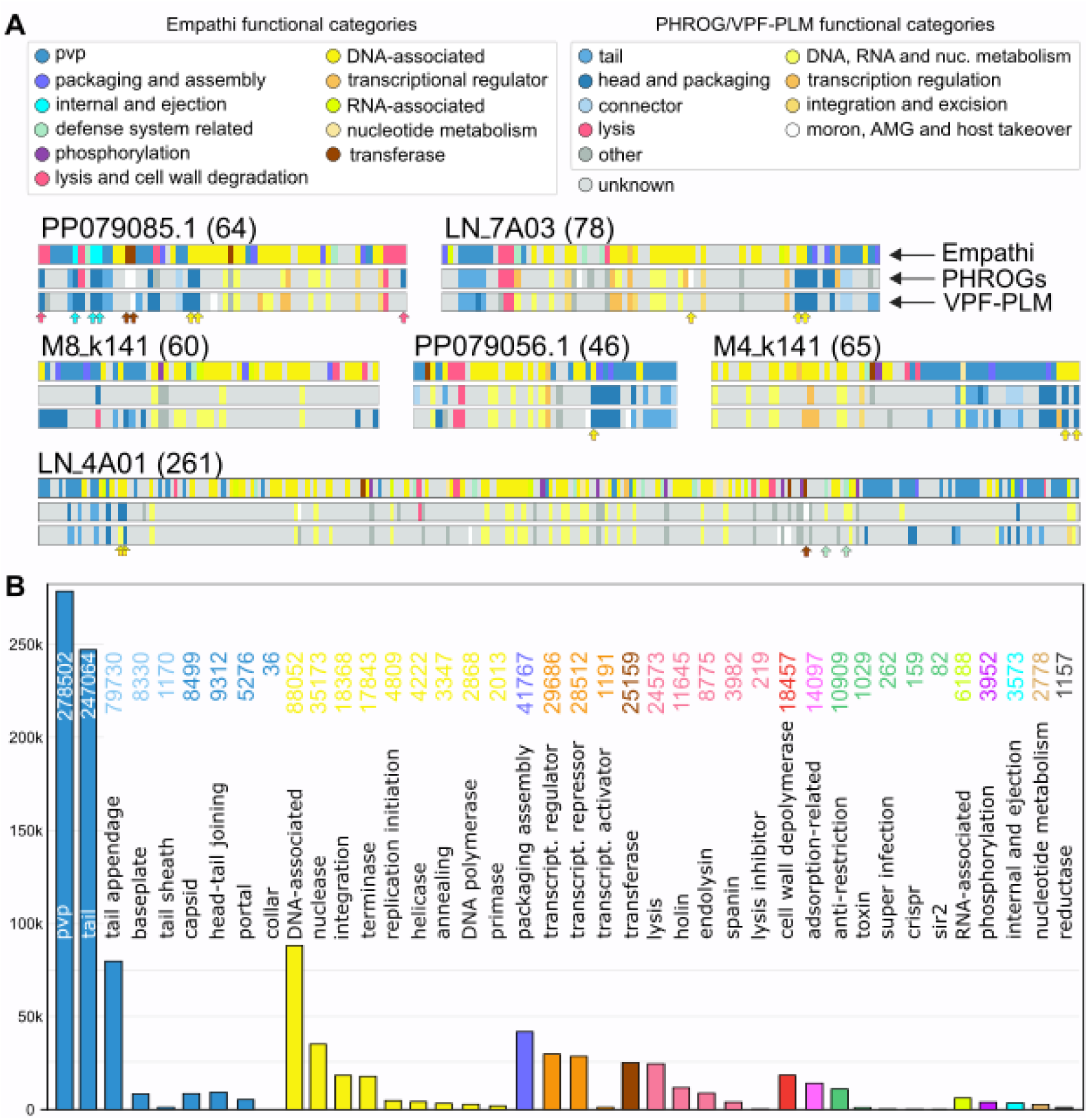
Empathi predictions on complete genomes and on the EnVhog database of phage proteins. **A)** Genomic maps of six genomes assembled from three viromes (Garmaeva et al.^39^, Bi et al.^40^, and Ni et al.^41^) coloured according to the functional annotations obtained using Empathi, pHMM comparison with the PHROG database and VPF-PLM^28^. Note that similar colours were chosen for corresponding functional categories between Empathi and PHROGs, and that each protein is represented with a uniform width in the genomic maps. Arrows indicate proteins for which the Empathi prediction correponds to the PHROG annotation term rather than category. The number of proteins found in each genome is indicated in parentheses. pvp: phage virion protein. PP079085.1 and PP079056.1 come from Ni et al., LN_4A01 and LN_7A03 come from Garmaeva et al., and M4_k141 and M8_k141 come from Bi et al. **B)** Empathi predictions for the EnVhog database: predicted protein counts for each functional group at a confidence > 0.95.

A more in-depth analysis of the predictions made for one genome (PP079085.1) from Ni *et al*.’s study (Figure 4) helps elucidate some of the observed differences (Supplementary table 2). First, two proteins predicted as being “lysis-associated proteins” (see red arrows) by Empathi are classified under the “head and packaging” category by PHROG pHMMs (phrog_2860). The PHROG *annotation terms* instead of *categories* reveal these proteins are indeed endolysins. Furthermore, VPF-PLM also predicted them as being “head and packaging” proteins which is expected from a model trained on the PHROG categories, yet the most important annotation – the one related to the actual function of the protein – is lost. Second, there are three proteins predicted by Empathi as being “internal/ejection proteins” (in cyan) and these are indeed internal proteins placed under the “head and packaging” category using PHROG pHMMs (phrog_308, phrog_418, phrog_6651). Third, two proteins predicted as “DNA-associated” and “packaging-related” (in yellow) by Empathi are a terminase small and large subunit which is also consistent (also in the “head and packaging” PHROG category; phrog_2, phrog_11494). Finally, two glycosyltransferases (see brown arrows) are predicted as being “transferases” by Empathi, but placed in the “moron, auxiliary metabolic gene and host takeover” PHROG category (phrog_14945, phrog_34859). Once again for these last three examples, VPF-PLM usually predicted the large category like with PHROG pHMMs but loses critical information about the actual molecular function of those proteins.

Of note, similar functions assigned by Empathi seem to be highly colocalized. Phage virion proteins, DNA-associated proteins and lysis-associated proteins are usually grouped together. This is consistent with the fact that proteins having similar functions are usually colocalized in phage genomes in order to be expressed at the same time.

### 3.3 Increasing the proportion of annotated orthologous groups in the EnVhog database

Using Empathi, the fraction of annotated viral orthologous groups in the EnVhog database is greatly increased from only 16% to 67.5% if we consider proteins that received a prediction with a confidence greater than the default threshold of 50%. The EnVhog database being composed of proteins obtained from metagenomic data, it is representative of a much greater diversity than the proteins used to train Empathi. Since proteins that have a more distant homology to proteins in the training set are more likely to lead to erroneous predictions, a more stringent confidence threshold of 95% was used to keep predictions, resulting in 33% of annotated protein clusters in the whole dataset, corresponding to a 17% increase compared to previous annotations. These can be visualized in Figure 4B. Once again, a great diversity of protein functions can be observed in the dataset, with phage virion proteins (mostly tail proteins) and DNA-associated proteins being the most abundant.

Finally, a random sample of 1000 proteins was taken from the previously unannotated fraction of the EnVhog database with the objective of comparing Empathi’s and VPF-PLM’s ability to assign predictions to new proteins. Empathi annotates 25.8% of proteins at a confidence of 95% (63.4% using default 50% confidence), while VPF-PLM can only annotate 17.3% using its default confidences (see calibrated thresholds in Flamholz *et al.*^28^).

## 4 Discussion

In this work we developed Empathi, a tool that leverages the highly informative representations generated by protein language models to annotate beyond standard and remote homology. It constitutes a significant improvement from the most recent model proposed by Flamholz *et al.* for this task. By using binary models and reorganizing annotations to make them more consistent with the molecular functions of proteins, we are able to improve the accuracy and sensitivity of models trained to predict protein functions. Beyond experimental validation using the testing dataset, the high colocalization of predicted functions in complete genomes further demonstrates Empathi’s consistency. Finally, Empathi was applied to the EnVhog database of phage proteins, doubling the proportion of annotated protein clusters from 16% to 33%.

Being constituted of independent binary models, Empathi can assign multiple functions to proteins such as a specific and a general function like “nuclease” and “DNA-associated”, or even multiple general functions. For example, some packaging proteins can also bind to DNA^44^, and some structural proteins can also have lytic domains^19^. Tools only assigning a single annotation strongly limit subsequent analyses. Out of the 15.4k predicted cell wall depolymerases – including lysins and exopolysaccharide (EPS) depolymerases – in the EnVhog database, 3k are also predicted as being structural proteins. This is important as it hints to the biological role of these proteins. EPS depolymerases and *virion associated* (structural) lysins intervene at the beginning of the infection process to enable the phage to insert its DNA into the host bacterium^19,45,46^ (“cell wall depolymerase” category in figure 1). Endolysins have a completely different function, serving to degrade the bacterial cell wall rapidly at the end of the infection process to liberate the newly produced phage virions (in “lysis” category in figure 1).

From a machine learning perspective, it is imperative that the functional groups we want to predict are consistent with the underlying biology. The PHROG categories were not originally intended as labels for ML; they contain enormous overlap that certainly adds noise and likely impacts the performances of models trained on them. For example, DNA-associated proteins are included in 7 of the 9 PHROG categories: 1) *DNA, RNA and nucleotide metabolism*, 2) *integration and excision*, 3) *transcription regulation*, 4) *moron, auxiliary metabolic gene and host takeover*, 5) *head and packaging*, 6) *tail* and 7) *other*. This means that there are proteins with very similar functions (in the DNA-associated category) that are present in the positive and in the negative sets for all of these classes when using the PHROG categories as training labels.

In addition, very little biological insights are gained by predicting that a protein belongs to the “other” category as is done in VPF-PLM because 1) the specific annotation that is present in the PHROG database is lost and 2) this category is simply an agglomeration of proteins with differing molecular functions. To name only a few, it is composed of methyltransferases, proteases, recombinases, kinases, lipoproteins, etc. The same can be said about the “moron, AMG and host takeover category” being composed of proteins with annotations such as “oxygenase”, “exclusion protein”, “toxin”, “glucosyltransferase”, “ABC transporter”, “ribonucleotide reductase”, etc.

Here, a reorganization of the PHROG categories into groups that share a common molecular function was realized in order to increase the accuracy, confidence and recall of models. In many cases, this required creating a hierarchy that can be used to differentiate proteins that share a common high-level function but that perform different tasks (lower-level functions). This is the case for structural proteins (capsid, tail, collar, etc.), for proteins associated to lysis at the end of the infection cycle (lysin, holin, spanin, lysis inhibition), and for DNA-associated proteins (nuclease, integrase, transcriptional regulation, etc.).

Even though it is difficult to evaluate the generalization capabilities of such a model, we evaluated how our tool would behave on non-homologous proteins by testing it on proteins with no detectable similarities to proteins in the training set (see holdout procedure in section 2.5). Empathi still performed well showing that even without any detectable similarities at the sequence level, protein embeddings still encode a signal that can be used to make predictions. Of course, as more proteins are discovered and added to databases, our model will likely need to be retrained to consider this new diversity.

Furthermore, the dataset used to train Empathi was annotated using PHROG pHMMs. This is currently the most well adapted method as it can provide the specific (low-level) annotations required to train our models, but they still remain secondary annotations not verified experimentally that could potentially introduce biases in our models. Having a large diversity of protein sequences in each group helps to mitigate these biases. Removing clusters containing proteins with differing annotations from the training set also reduces potential noise. Still, a feature of our tool is to provide the confidence of each prediction. With this information, a user can choose the desired trade-off between high specificity (confidence threshold of 95-99%) and high sensitivity (default threshold of 50%).

We believe this tool will provide a more comprehensive view of the functions possessed by phages helping to better characterize and understand them.

## Data and method availability

Empathi software can be freely downloaded at https://huggingface.co/AlexandreBoulay/EmPATHi and the information pertaining to the data used to train models or for supplementary analyses can be found at https://doi.org/10.5281/zenodo.14036012.

