## Supplementary figure 1 for "Empathi: Embedding-based Phage Protein Annotation Tool by Hierarchical Assignment"

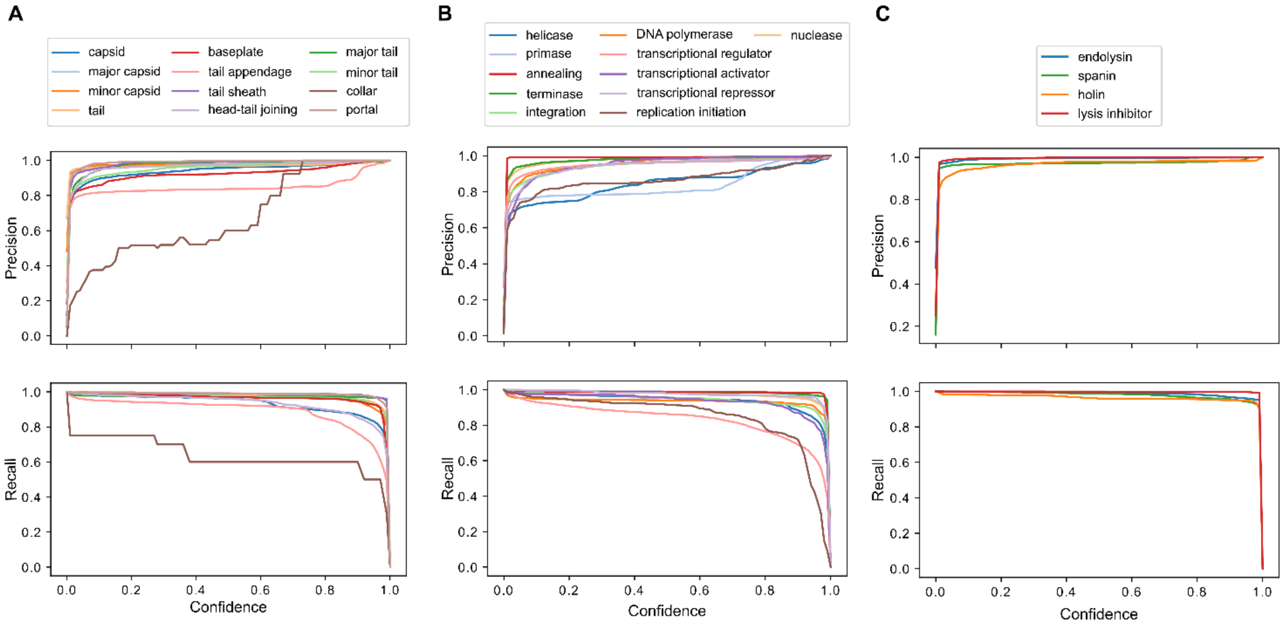


***Supplementary figure 1****. Precision and recall curves as a function of confidence for Empathi models trained on* ***A)*** *subcategories of structural proteins,* ***B)*** *subcategories of DNA-associated proteins and* ***C)*** *subcategories of lysis-associated proteins.*
