## Supplementary table 1 for "Empathi: Embedding-based Phage Protein Annotation Tool by Hierarchical Assignment"

*Supplementary table 1. Number of proteins and protein clusters used to train and test Empathi models as well as performance metrics obtained on the test set.*

| | Training | | | | Testing | | | | Empathi testing scores | | |
|---|---|---|---|---|---|---|---|---|---|---|---|
| | # Pos clusters | # Pos proteins | # Neg clusters | # Neg proteins | # Pos clusters | # Pos proteins | # Neg clusters | # Neg proteins | F1-score | Precision | Recall |
| **Packaging and assembly** | 2723 | 26373 | 36668 | 36668 | 666 | 6338 | 9182 | 9182 | 0.97 | 0.98 | 0.95 |
| **PVP** | 12971 | 108029 | 26404 | 26404 | 3219 | 28080 | 6625 | 6625 | 0.98 | 0.99 | 0.96 |
| **Tail** | 8304 | 67259 | 2640 | 28392 | 2086 | 15886 | 650 | 7422 | 0.98 | 0.97 | 0.98 |
| **Major tail** | 428 | 5324 | 6373 | 53817 | 94 | 647 | 1607 | 11002 | 0.98 | 0.99 | 0.97 |
| **Minor tail** | 1363 | 16107 | 5408 | 39877 | 346 | 4260 | 1347 | 9507 | 0.97 | 0.97 | 0.98 |
| **Capsid** | 1219 | 12471 | 9734 | 82447 | 282 | 2828 | 2457 | 21446 | 0.96 | 0.96 | 0.96 |
| **Major capsid** | 510 | 7616 | 559 | 4090 | 145 | 1625 | 123 | 805 | 0.98 | 0.98 | 0.99 |
| **Minor capsid** | 487 | 3337 | 581 | 8597 | 118 | 1054 | 150 | 1126 | 0.98 | 0.98 | 0.97 |
| **Baseplate** | 964 | 11108 | 5825 | 46684 | 251 | 2403 | 1447 | 10542 | 0.95 | 0.93 | 0.97 |
| **Tail appendage** | 2647 | 12798 | 3959 | 41375 | 644 | 3639 | 1008 | 10989 | 0.88 | 0.84 | 0.92 |
| **Portal** | 520 | 6711 | 10458 | 85116 | 130 | 1669 | 2615 | 26237 | 0.99 | 1.00 | 0.99 |
| **Collar** | 31 | 213 | 10943 | 95230 | 8 | 20 | 2736 | 24161 | 0.60 | 0.60 | 0.60 |
| **Tail sheath** | 222 | 2458 | 6579 | 54462 | 59 | 851 | 1642 | 13029 | 0.99 | 1.00 | 0.99 |
| **Head-tail joining** | 717 | 8860 | 10257 | 89311 | 173 | 1397 | 2571 | 20126 | 0.97 | 0.98 | 0.96 |
| **DNA-associated** | 12581 | 105873 | 24857 | 24857 | 3106 | 27216 | 6254 | 6254 | 0.98 | 0.99 | 0.96 |
| **Integration** | 1792 | 14657 | 9876 | 84294 | 465 | 3192 | 2453 | 23266 | 0.96 | 0.97 | 0.96 |
| **Nuclease** | 3406 | 27434 | 8254 | 72908 | 857 | 6890 | 2059 | 18200 | 0.98 | 0.99 | 0.99 |
| **DNA polymerase** | 717 | 8799 | 10843 | 90200 | 180 | 1817 | 2710 | 23414 | 0.96 | 0.98 | 0.94 |
| **Terminase** | 1124 | 11456 | 10569 | 88568 | 255 | 3374 | 2669 | 23646 | 0.99 | 0.99 | 0.99 |
| **Annealing** | 275 | 2757 | 11431 | 97824 | 68 | 877 | 2859 | 25471 | 0.99 | 0.99 | 0.99 |
| **Helicase** | 811 | 9913 | 10714 | 90371 | 182 | 1579 | 2700 | 21902 | 0.91 | 0.87 | 0.96 |
| **Primase** | 345 | 3584 | 11215 | 94299 | 69 | 1182 | 2822 | 25432 | 0.88 | 0.80 | 0.99 |
| **Replication initiation** | 395 | 1642 | 11289 | 100722 | 83 | 377 | 2839 | 24146 | 0.89 | 0.86 | 0.92 |
| **Transcriptional regulator** | 3183 | 14695 | 35429 | 35429 | 776 | 3307 | 8877 | 8877 | 0.91 | 0.96 | 0.86 |
| **Transcriptional activator** | 158 | 924 | 1530 | 7401 | 40 | 209 | 382 | 1696 | 0.97 | 0.99 | 0.95 |
| **Transcriptional repressor** | 842 | 3057 | 846 | 5521 | 200 | 515 | 222 | 1137 | 0.97 | 0.97 | 0.98 |
| **Adsorption-related** | 2581 | 12561 | 36703 | 36703 | 671 | 3558 | 9150 | 9150 | 0.93 | 0.92 | 0.94 |
| **RNA-associated** | 892 | 9347 | 38580 | 38580 | 257 | 2820 | 9612 | 9612 | 0.94 | 0.99 | 0.90 |
| **Nucleotide metabolism** | 920 | 12696 | 38556 | 38556 | 240 | 2898 | 9629 | 9629 | 0.98 | 0.99 | 0.98 |
| **Phosphorylation** | 1057 | 11182 | 38408 | 38408 | 245 | 1676 | 9622 | 9622 | 0.95 | 0.97 | 0.94 |
| **Transferase** | 1621 | 13721 | 37846 | 37846 | 385 | 2517 | 9482 | 9482 | 0.96 | 0.98 | 0.94 |
| **Reductase** | 455 | 6881 | 39043 | 39043 | 121 | 1851 | 9754 | 9754 | 0.98 | 1.00 | 0.97 |
| **Ejection** | 438 | 4364 | 38265 | 38265 | 99 | 655 | 9577 | 9577 | 0.93 | 0.94 | 0.92 |
| **Cell wall depolymerase** | 2038 | 19677 | 37373 | 37373 | 482 | 4003 | 9371 | 9371 | 0.93 | 0.98 | 0.89 |
| **Anti-restriction** | 964 | 6580 | 38503 | 38503 | 228 | 1825 | 9639 | 9639 | 0.96 | 0.99 | 0.93 |
| **Crispr** | 44 | 269 | 39453 | 39453 | 6 | 149 | 9869 | 9869 | 0.98 | 0.98 | 0.97 |
| **Sir2** | 39 | 549 | 39461 | 39461 | 4 | 39 | 9872 | 9872 | 1.00 | 1.00 | 1.00 |
| **Super infection** | 163 | 817 | 39315 | 39315 | 45 | 376 | 9825 | 9825 | 0.95 | 0.96 | 0.94 |
| **Toxin** | 271 | 1013 | 39189 | 39189 | 59 | 280 | 9806 | 9806 | 0.90 | 0.98 | 0.83 |
| **Lysis** | 2030 | 18289 | 37418 | 37418 | 469 | 5286 | 9394 | 9394 | 0.96 | 0.98 | 0.94 |
| **Endolysin** | 895 | 9140 | 1102 | 9239 | 242 | 2412 | 258 | 2658 | 0.99 | 1.00 | 0.99 |
| **Holin** | 561 | 4163 | 1436 | 14453 | 143 | 1396 | 357 | 3437 | 0.97 | 0.98 | 0.96 |
| **Spanin** | 383 | 2923 | 1616 | 15422 | 111 | 839 | 389 | 4391 | 0.98 | 0.97 | 0.99 |
| **Lysis inhibitor** | 146 | 1364 | 1853 | 17038 | 44 | 1290 | 456 | 3883 | 1.00 | 1.00 | 1.00 |
