## Supplementary table 2 for "Empathi: Embedding-based Phage Protein Annotation Tool by Hierarchical Assignment"

Supplementary table 2. Proteins in the PP079085.1 genome from Ni et al. study for which the Empathi prediction differs from the PHROG category but is coherent with the more precise PHROG annotation.

| Position in genome | Empathi | PHROG annotation | PHROG category | PHROG entry | VPF-PLM |
|---|---|---|---|---|---|
| 1 | lysis | endolysin | Head and packaging | phrog_2860 | Head and packaging |
| 8 | Internal/ejection | Internal virion protein | Head and packaging | phrog_308 | Head and packaging |
| 11 | Internal/ejection | Internal virion protein | Head and packaging | phrog_418 | Head and packaging |
| 12 | Internal/ejection | Internal virion protein | Head and packaging | phrog_6651 | Head and packaging |
| 17 | Transferase | glycosyltransferase | moron, auxiliary metabolic gene and host takeover | phrog_2 | Head and packaging |
| 18 | Transferase | glycosyltransferase | moron, auxiliary metabolic gene and host takeover | phrog_11494 | Head and packaging |
| 28 | DNA-associated Packaging/assembly | Terminase large subunit | Head and packaging | phrog_14945 | Unknown |
| 29 | DNA-associated Packaging/assembly | Terminase small subunit | Head and packaging | phrog_34859 | Moron, AMG and host takeover |
| 46 | lysis | endolysin | Head and packaging | phrog_2860 | Unknown |
